# Changes in Iris Stiffness and Permeability in Primary Angle Closure Glaucoma

**DOI:** 10.1101/2021.05.04.442554

**Authors:** Satish K. Panda, Royston K. Y. Tan, Tin A. Tun, Martin L. Buist, Monisha Nongpiur, Mani Baskaran, Tin Aung, Michaël J. A. Girard

## Abstract

**Purpose:** To evaluate the biomechanical properties of the iris by evaluating iris movement during pupil constriction and to compare such properties between healthy and primary angle-closure glaucoma (PACG) subjects.

**Methods:** A total of 140 subjects were recruited for this study. In a dark room, the anterior segments of one eye per subject were scanned using anterior segment optical coherence tomography (AS-OCT, SS-1000 CASIA, Tomey Corporation, Nagoya, Japan) imaging during induced pupil constriction with an external white light source of 1700 lux. Using a custom segmentation code, we automatically isolated the iris segments from the AS-OCT images, which were then discretized and transformed into a three-dimensional point cloud. For each iris, a finite element (FE) mesh was constructed from the point cloud, and an inverse FE simulation was performed to match the clinically observed iris constriction in the AS-OCT images. Through this optimization process, we were able to identify the elastic modulus and permeability of each iris.

**Results:** For all 140 subjects (95 healthy and 45 PACG of Indian/Chinese ethnicity, Age: 60.2±8.7 for PACG subjects and 57.7±10.1 for healthy subjects), the simulated deformation pattern of the iris during pupil constriction matched well with OCT images. We found that the iris stiffness was higher in PACG than in healthy controls (24.5±8.4 kPa vs 17.1±6.6 kPa with 40 kPa of active stress specified in the sphincter region; *p* < 0.001), whereas iris permeability was lower (0.41±0.2 mm^2^/kPa.s vs 0.55±0.2 mm^2^/kPa.s; *p* = 0.142).

**Conclusion:** This study suggests that the biomechanical properties of the iris in PACG are different from those in healthy controls. An improved understanding of the biomechanical behavior of the iris may have implications for the understanding and management of angle-closure glaucoma.

## Introduction

Primary angle-closure glaucoma (PACG) causes irreversible blindness affecting four million people worldwide [1]. A narrower anterior chamber angle is the main predisposing characteristic of eyes suffering PACG [2, 3]. Several anatomical features, such as a shallow anterior chamber depth, a short axial length, and a small anterior chamber width, are the other major PACG risk factors [4]. However, several studies have reported that only a small proportion of those with narrow angles or with other PACG risk factors develop PACG [5, 6]. In the study by Friedman et al., PACG eyes responded differently to changes in illumination compared to healthy eyes with the same level of narrow angles [7]. This suggests that a static morphologic assessment of the anterior chamber may not be sufficient PACG diagnosis, and instead, dynamic physiologic changes of ocular structures (such as iris movement during miosis/mydriasis) could prove pertinent.

Several studies have reported associations between iris biomechanics and PACG. For instance, the density of collagen fibers in the iris is higher in PACG than in healthy subjects as characterized through histology [4]. Narayanaswamy et al. performed biomechanical testing on excised iris strips and found that PACG eyes exhibited a higher iris stiffness compared to healthy eyes [8]. However, characterizing the stiffness of the iris in an ex-vivo setting has poor clinical applicability because it requires an invasive procedure to harvest a large biopsy specimen. Because of this limitation, Pant et al. proposed a non-invasive in-vivo procedure to evaluate iris stiffness directly from optical coherence tomography (OCT) images [9]. While this was an important piece of work, it was limited by the assumption that the iris was an incompressible solid that could not exchange water with the anterior chamber. Because of this limitation, the reported stiffness measurements may be inaccurate.

From a biomechanical point of view, the iris is composed of two phases, a solid phase and a fluid phase [10]. It contains a large amount of water that can easily flow in and out during miosis or mydriasis. The ability of the iris to absorb or exude fluid is defined by its permeability, with a high permeability allowing more water movement across the iris. This water movement can affect the entire volume of the iris. For instance, several studies have shown that the volume of the iris can change during pupillary constriction/dilation [11]. In addition, iris surface features, known as crypts and furrows, were found to be correlated with the rate of volume change [11, 12, 13]. It has also been reported that eyes with thicker and larger irides (i.e., a larger iris volume) are more prone to angle-closure [14]. Thus, measuring the permeability of the iris in-vivo (in addition to its stiffness) may provide useful information about its pathophysiological states.

In this study, we aimed to evaluate the stiffness and permeability of the iris during pupil constriction using an inverse finite element (FE) approach in both PACG and age-matched healthy subjects. We considered the iris to be a heterogeneous structure with a permeable stroma layer and modeled it as a biphasic material thus allowing for water movement.

## Methods

### Patient recruitment and imaging

A total of 140 subjects (95 healthy and 45 PACG) were recruited for this study at the Singapore Eye Research Institute, Singapore. Written informed consent was obtained for all patients. The study was conducted following the tenets of the World Medical Associations Declaration of Helsinki and had ethics approval from the SingHealth Centralized Institutional Review Board. All subjects underwent anterior segment imaging using swept-source OCT imaging (SS-1000 CASIA, Tomey Corporation, Nagoya, Japan) in the primary gaze position before any contact procedures. OCT imaging was performed with the “angle analysis” protocol (video mode) to obtain ‘live’ horizontal B-Scans of the anterior segment (0^0^ to 180^0^, limbal to limbal), and the acquisition time was 0.125 second per line, i.e., eight frames per second. The recording of the OCT video was started one minute after dark adaption using a standard protocol (light intensity was approximately 20 lux as measured by Studio Deluxe II L-398, Sekonic, Japan). A torchlight flashed the fellow eye from the temporal side 15^0^ off-axis (approximately 1700 lux) [37]. Changes in the anterior chamber and iris from dark to light were acquired. If motion artifacts were observed, the procedure was repeated up to a maximum of three times to prevent iris muscle fatigue. Each frame was 16 mm in length and 6 mm in depth.

### Segmentation of OCT images and 3D Reconstruction

We extracted ten frames from the OCT video of each subject such that the complete iris movement, from its initial fully-dilated position to its deformed position (fully-constricted), could be observed. The iris roots (IR) on both sides were identified in each frame, and the images were rotated in such a way that both IRs were aligned horizontally (see Fig. 1a and 1b). Using a custom Python code, we delineated the left and right iris cross-sectional boundaries in each of the ten frames manually and discretized them into a set of points (Figs. 1c-1e).

**Figure 1:**
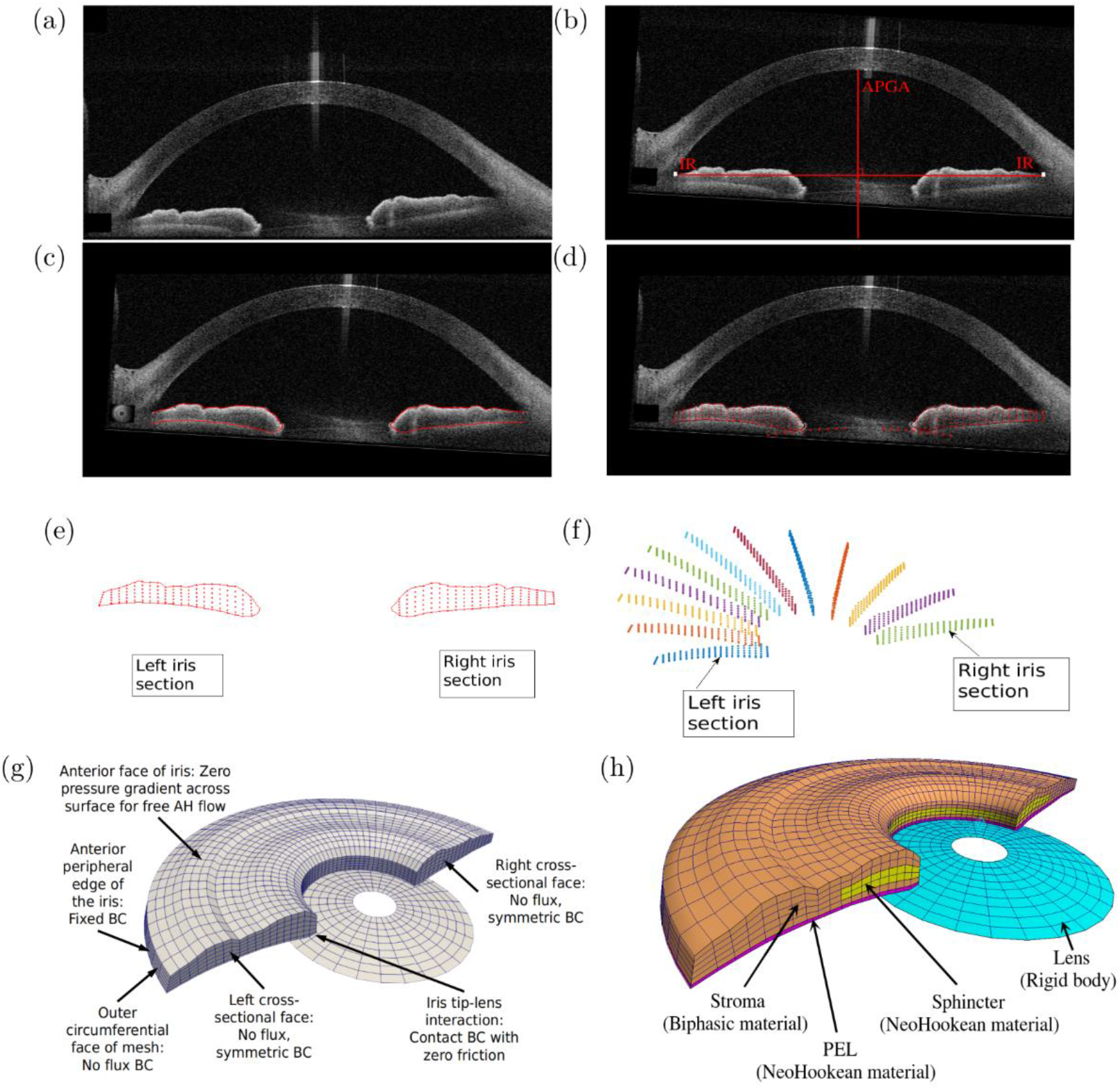
a) An OCT image of the iris and the anterior chamber of the eye. b) Rotated OCT image to align the IR horizontally. The IR and the APGA are shown with white dots and with a vertical red line, respectively. c-d) The OCT image with left and right iris boundaries marked and discretized into a set of points. e) The left and right iris boundaries extracted from the OCT image. f) Three-dimensional transformation and rotation of the left and right iris boundaries to generate a 3D point cloud. g) A 3D FE mesh of the iris generated from the point cloud with the specified boundary condition for FE simulation. h) Specified tissue layers (i.e., stroma layer in brown, pigment epithelial layer in pink, and sphincter layer in yellow) with the corresponding materials in the iris mesh.

We assumed the iris to be a rotationally symmetric structure around an anteroposterior geometric axis (APGA). The midpoint of the line segment joining both IRs was considered the origin, and the bisecting perpendicular line was treated as the APGA (Fig. 1b). The boundary points on the left cross-section of the iris were then rotated by 180^0^ about the APGA to create a three-dimensional (3D) point cloud (Fig. 1f). During the rotation, the points on the left face were transformed in such a way that the shape of the left iris would transform into that of the right after the rotation. Using a custom Python code, at first, we generated a node connectivity matrix and a 3D mesh for the iris, consisting of 4,600 eight-node trilinear hexahedral elements and the different tissue layers (i.e., stroma, iris pigment epithelial layer (PEL), and sphincter layer) were then identified in the mesh (Fig. 1g and 1h). The anterior boundary layer of the iris was considered to be a part of the stroma as it was thought to be unlikely to provide any significant mechanical resistance during deformation. Similarly, the dilator region was not specified as our study primarily focused on pupil constriction but not dilation. The PEL is a single-cell layer on the posterior side of the iris that restricts the movement of water. We thus created a thin PEL in our mesh to prevent any fluid movement from the posterior side. During miosis/mydriasis, the iris tip slides over the lens surface. We therefore digitized the boundary of the lens in the OCT image and meshed it with 4-noded linear quadrilateral shell elements (Fig. 1d and 1g). For simplicity, the lens was treated as a non-deformable material (also known as rigid body), over which the iris could slide during constriction.

### Constitutive relationships for all tissue layers

The iris was assumed to be a heterogeneous structure. Specifically, the sphincter and PEL were considered to be nearly incompressible and to behave as Neo-Hookean materials. Their constitutive relationships were defined as:

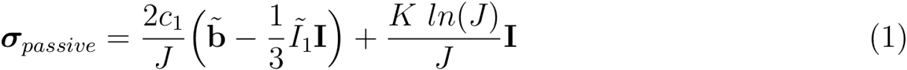

where σ_passive_ is the passive Cauchy stress tensor, c_1_ is the Neo-Hookean material coefficient, *Ĩ*_*1*_ is the invariant of the deviatoric part of the right Cauchy-Green deformation tensor defined as 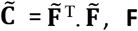 is the deformation gradient tensor and 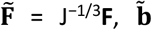 is the left Cauchy-Green deformation tensor defined as 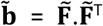, J is the Jacobian of the deformation gradient defined as J=det(**F**), *K* is a bulk modulus-like penalty parameter, and **I** is the second-order identity tensor. The Neo-Hookean material coefficient c_1_ is related to the elastic modulus of the material through E = 4c_1_(1 + ν), where ν is the Poisson’s ratio (ν was fixed at 0.499 for our simulations) [17].

The sphincter region was assumed to be electrically active, and the active force produced by the smooth muscle cells (SMCs) in the circumferential direction was defined as σ_active=_ σ e_θ_⊗e_θ_, where σ is the magnitude of the active force and e_θ_ is the unit circumferential vector. Following previous studies, the magnitude of σ_active_ was assumed to be 40 kPa for both PACG and healthy eyes [9, 18]. The total stress in the sphincter region was computed by adding the passive and active components, i.e., σ_total_ = σ_passive_ + σ_active_.

The stroma of the iris is a sponge-like structure that is composed of 40% water [16]. The aqueous humor (AH) in the anterior chamber is believed to readily flow in and out of the stroma during constriction and dilation of the pupil [14]. Although the cells of the stroma are not compressible, the flow of AH through the tissue boundary makes it compressible [15]. This phenomenon was observed in-vivo in previous studies [19, 20]. To capture this behavior, the stroma was considered to be a biphasic material consisting of a mixture of a porous-permeable solid and an interstitial fluid. Both the solid and fluid constituents of the mixture were assumed to be intrinsically incompressible. However, the mixture is compressible because the pores of the solid matrix may gain or lose fluid during the deformation of the mixture. The Cauchy stress for a biphasic material, σ_biphasic_ is given by:

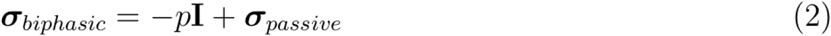

where *p* is the interstitial fluid pressure and σ_passive_ is the stress in the solid matrix. We assigned the same nearly incompressible material, as in Eq. 1, to describe the solid matrix of the stroma. The constitutive relation for the hydraulic permeability of the interstitial fluid flowing within the porous solid matrix was defined as:

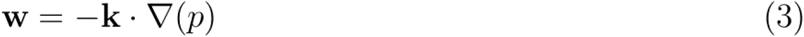

where **w** is the volumetric flux of the fluid relative to the solid, ∇(*p*) is the interstitial fluid pressure gradient, and **k** is the is the hydraulic permeability tensor. We also assumed the stroma exhibited a constant isotropic permeability, thus the hydraulic permeability tensor was defined as **k** = *k***I**, where *k* is simply referred as the iris permeability.

### Boundary conditions for all FE simulations

To recreate the physiological deformation pattern of the iris and associated AH flow during pupil constriction, we applied appropriate different boundary conditions to the FE models (Fig. 1g). For all FE simulations, we fixed the anterior peripheral edge of the iris mesh to mimic its connections with the trabecular meshwork. The nodes on the left/right cross-sectional faces were assigned a symmetric boundary condition, i.e., the out-of-plane motion was restricted. We also defined no-flux boundary conditions for the outer circumferential face and the left/right cross-sectional faces of the mesh to prevent AH flow through these faces. However, the AH was allowed to flow through the anterior surface of the iris during constriction. Thus, the pressure difference across this surface was set to p=0 Pa. Note that during pupil constriction, the iris tip slides over the lens surface. To mimic this phenomenon, we defined a sliding-contact boundary condition between the iris tip and the lens surface with a coefficient of friction equal to zero. No external loads were applied to the iris, and its deformations were solely due to the active force produced by the sphincter during contraction. Each FE model was solved using FEBio (3.0, Musculoskeletal Research Laboratories, University of Utah, UT, USA) – a nonlinear FE solver designed for biomechanical studies [21].

### Extraction of Iris Biomechanical Properties

Each iris FE model was characterized by two material parameters, the elastic modulus, c_1_ and the permeability, *k*. To extract these parameters for each iris, we used an inverse FE approach that aimed to match the motion of the iris in the FE simulations to that observed in the OCT images. Specifically, the active force of the sphincter muscle was increased in ten incremental steps, and in each step, the parameters were optimized so that the cross-sectional shape of the iris in the FE simulation best-matched that observed with OCT imaging. For this optimization process, the cost function to be minimized was defined as:

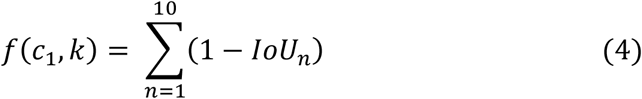

where IoU_n_ is the intersection over the union of iris cross-sections (FE vs OCT) for a given time step, n. The IoU is a commonly used metric for measuring overlap between two shapes. The IoU is equal to 1 for two shapes that are perfectly overlapped, and equal to zero when there is no overlap. To obtain the biomechanical properties of each iris, Equation (4) was minimized using the Nelder-Mead algorithm, which is a heuristic search method that can converge to a local minimum [22]. The optimization procedure was therefore repeated multiple times with different initial guesses to ensure global convergence.

A C++ based framework coupled with Unfit (Computational Bioengineering lab, NUS) was developed to automate the whole process. Unfit is a C++ based nonlinear optimization software suitable for data fitting problems. This framework called the FEBio solver during each iteration of the optimization and then minimized the objective function. Unfit also provided options to define bounds for the variables while using the Nelder-Mead algorithm. We defined physically realistic bounds for the iris of 0.001-50 kPa for c_1_ and 0.001-10 mm^2^/kPa.s for k.

To examine the effects of the active stress in the sphincter region on the extracted stiffness and permeability values, we varied the active stress from 20 kPa to 50 kPa in steps of 5 kPa and performed an optimization at each stress level. The optimized stiffness and permeability values were plotted against the applied active stress, and Pearson’s correlation (PC) tests were performed. The PC tests provide an output that lies between -1 and 1, where 0 indicates no correlation, and a value below -0.5 or above 0.5 indicates a notable correlation [23].

## Results

The subjects in our study were of Indian/Chinese ethnicity with an average age of 60.2±8.7 for PACG subjects and 57.7±10.1 for healthy subjects.

We found the iris to be stiffer in PACG eyes compared to healthy eyes (*p* < 0.001, Fig. 2a), whereas the permeability was found to be lower (*p* = 0.142, Fig. 2b) in PACG eyes. The elastic moduli of healthy and PACG irides with 40 kPa active stress were 17.1±6.6 kPa and 24.5±8.4 kPa, respectively; whereas the respective permeability values were 0.55±0.2 mm^2^/kPa.s and 0.41±0.2 mm^2^/kPa.s. The tissue stiffness was found to increase proportionally with the applied active stress (PC = 0.974, Fig. 2c), whereas the permeability value was uncorrelated with the active stress (PC = -0.193, Fig. 2d).

**Figure 2:**
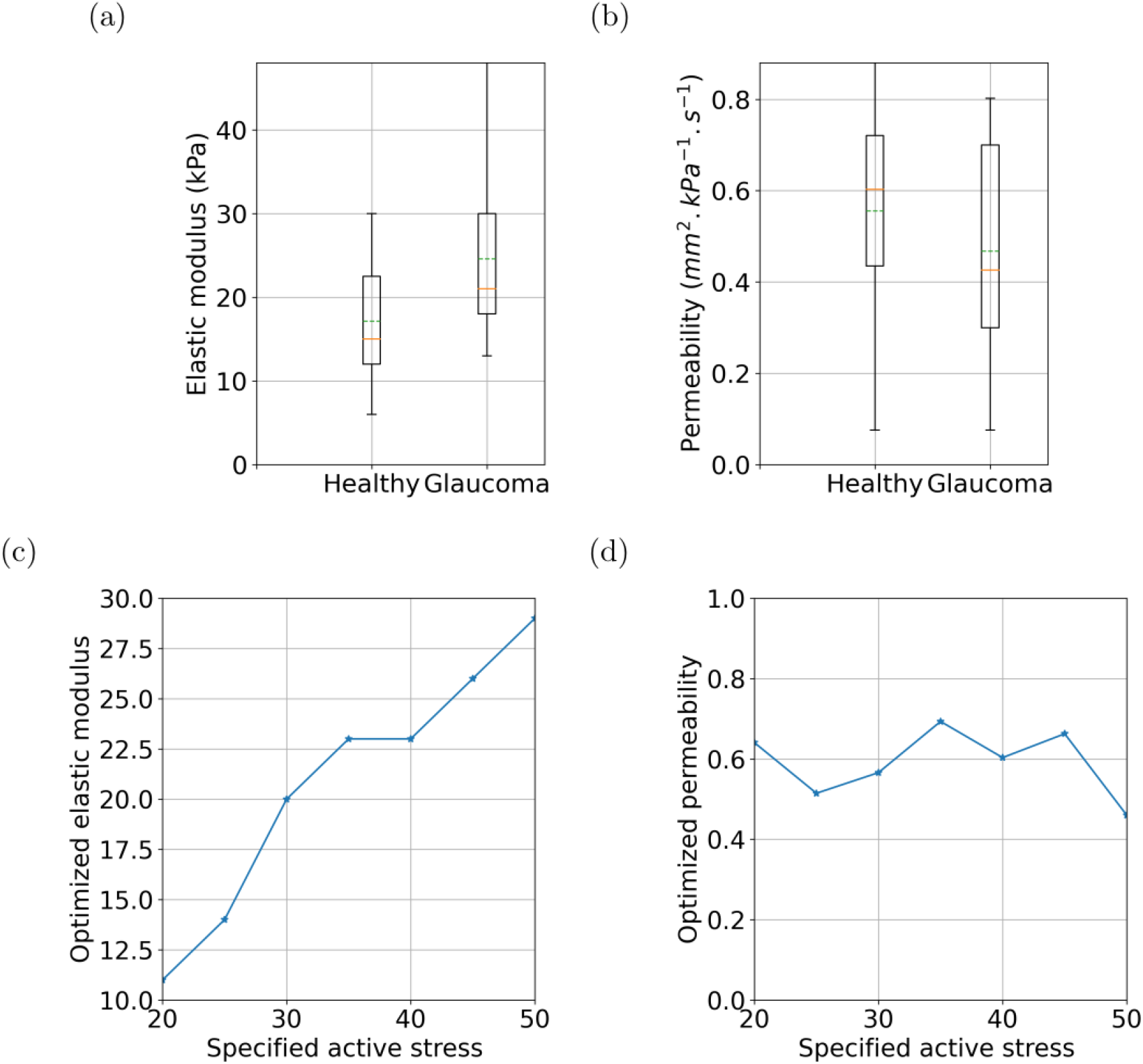
Comparison of a) stiffness and b) permeability values of healthy and PACG irides with 40 kPa of active stress in the sphincter region. c,d) Correlation of tissue stiffness and permeability with the specified active stress.

For all incremental loading steps, our FE simulation results followed the actual iris motion closely (IoU > 0.82 for all eyes). Fig. 3 shows the iris deformations for one subject at six different time steps with the ‘true’ iris boundary in red and the FE-simulated iris boundary in blue. Overall, this suggests that our proposed optimization framework was successful in matching the observed motion of the iris.

**Figure 3:**
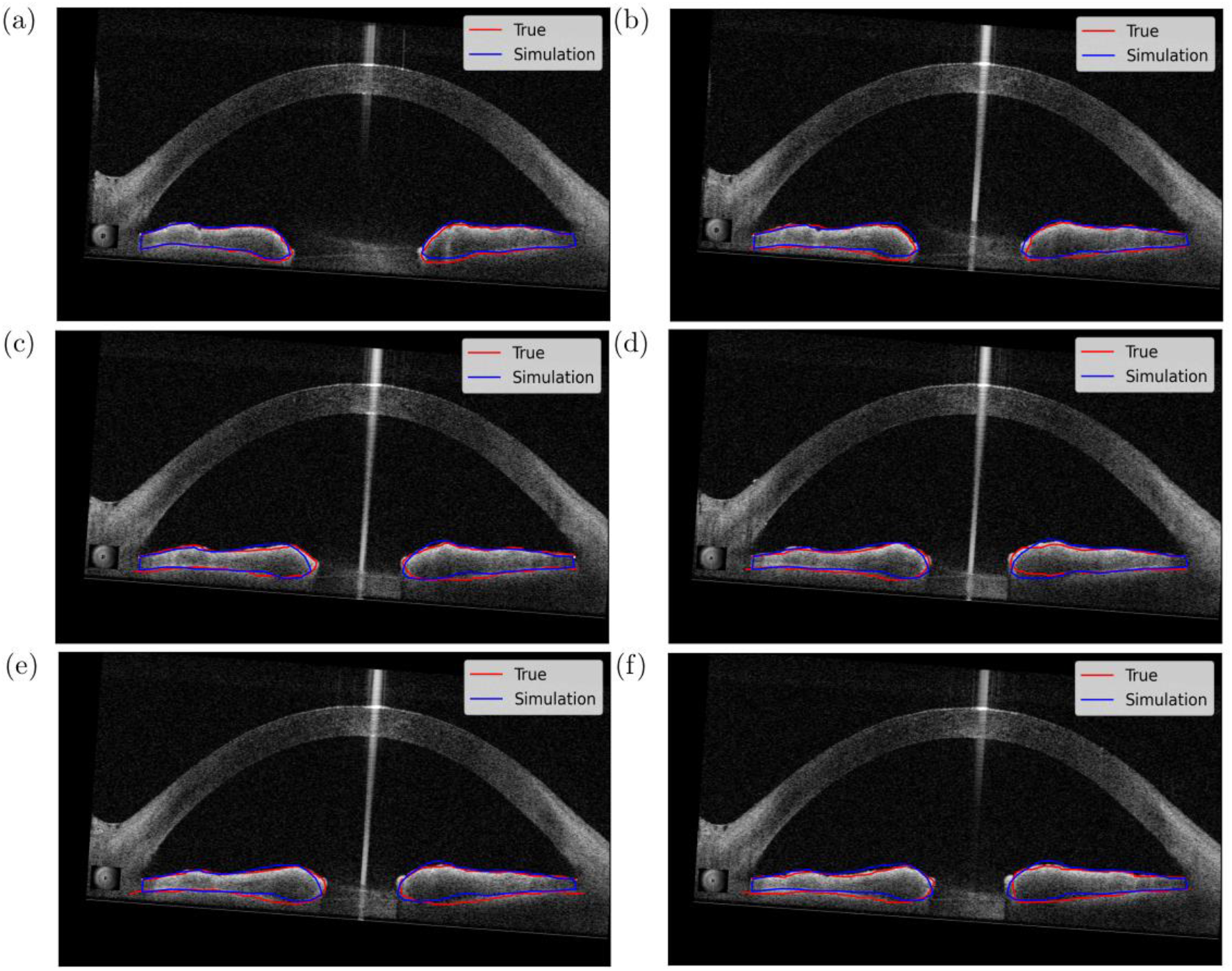
a-f) Deformation of the iris at different instants of time during pupil constriction. The red lines are the actual boundaries of the iris, whereas the blue lines are the FE simulation results.

Fig. 4a depicts the iris mesh in its initial undeformed configuration in our FE simulation, and Fig. 4b shows the specified active stress in the sphincter region. As a consequence of the active stress, the iris mesh deformed, and Fig. 4c shows the magnitude of strain at the peak of the deformation. We noted that during iris constriction, the AH flowed into to the iris through the anterior face of the stroma, whereas it flowed out near the iris root and tip (see Fig. 4d). This behavior was consistent across all eyes.

**Figure 4:**
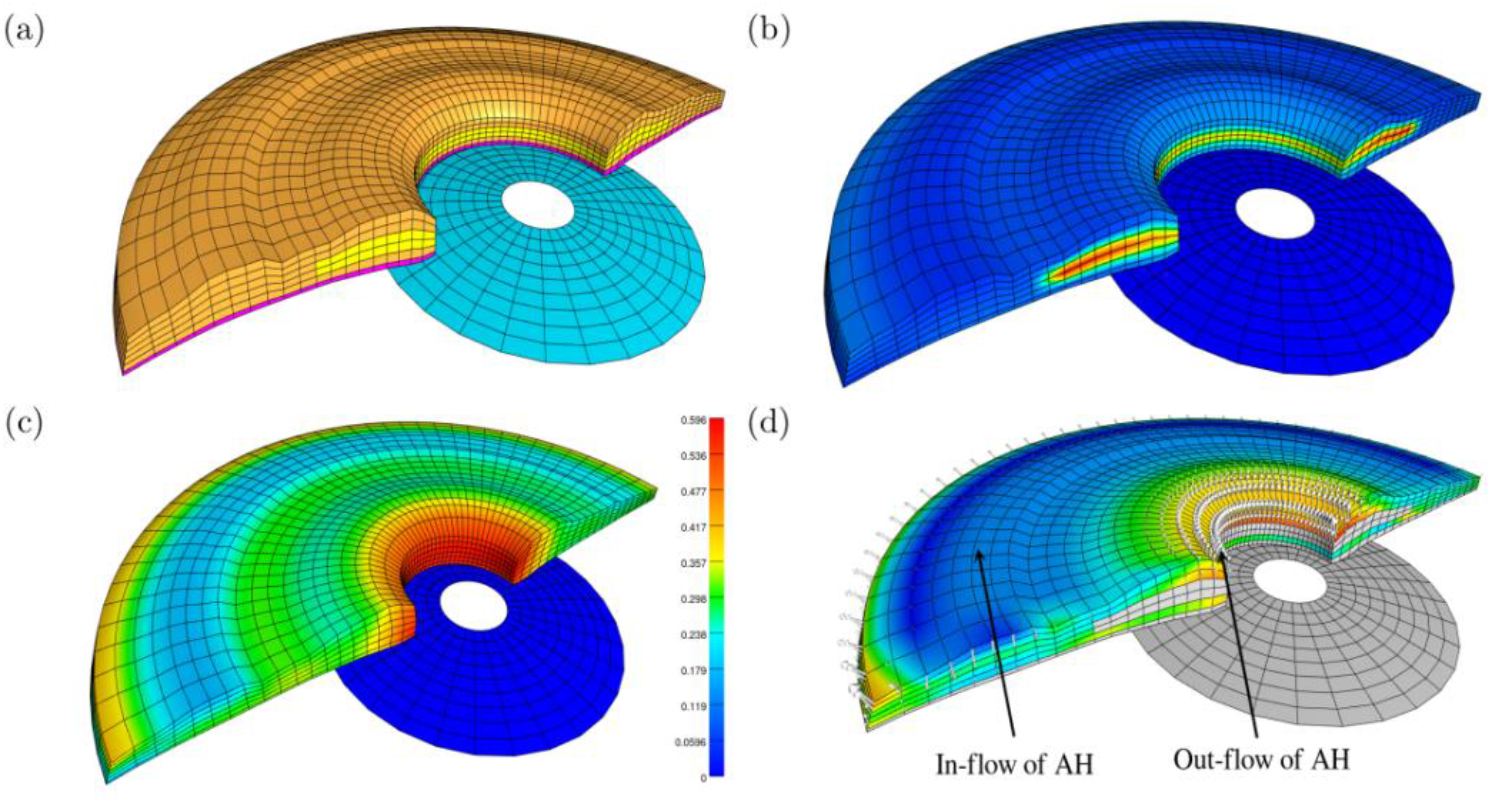
a) Iris mesh in its undeformed reference configuration. b) Applied active stress in the sphincter region. c) Iris deformation because of the active stress in the sphincter region. Color code shows the magnitude of strain at the peak of constriction. d) Flow of aqueous humor through the stroma at the peak of the constriction.

The mesh convergence study showed that changes in the objective function did not change beyond a total number of 4,600 elements. Therefore, the same number of elements were used for all FE simulations.

## Discussion

In this study, we used an inverse FE method to evaluate the biomechanical properties of the iris stroma in both PACG and healthy subjects. Our FE model was able to replicate the iris motion observed clinically with OCT imaging during pupil constriction. Through this process we were able to extract the stiffness and permeability of each iris. Overall, we found that in PACG subjects the iris stiffness was higher and permeability lower compared to healthy subjects.

### Higher tissue stiffness in PACG subjects

Our study revealed that the iris was stiffer in PACG subjects. This may be explained by the higher density of collagen fibers that has been observed in PACG irides. Indeed, several studies have reported that the density of Type-I collagen in the iris stroma was significantly higher in PACG eyes than in healthy eyes [4, 24]. For a given soft tissue, the density of collagen fibers typically correlates with the mechanical stiffness of such a tissue [25]. Using atomic force microscopy, Narayanaswamy et al. compared the elastic moduli of PACG and normal irides and reported values of 2.40±0.82 kPa and 0.85±0.31 kPa, respectively [8]. Pant et al. reported similar observations using an in-vivo and non-invasive inverse FE method [9]. Our results are consistent with these observations, but our methodology also had the benefit to additionally assess the permeability of the iris tissue.

The in-vivo stiffness of a healthy iris in our study was found to be 17.1±6.6 kPa with 40 kPa of applied active stress. In a recent study, Lee et al. estimated the murine iris stiffness to be 96.1±54.7 kPa following an in-vivo protocol [26]. The in-vivo iris stiffness of human subjects reported by Pant et al. was 38.8±15.8 kPa with 40 kPa of active stress. Furthermore, in a study by Heys and Barocas, the ex-vivo stiffness of bovine iris in the radial direction under uniaxial tensile testing was found to be 27.0±4.0 kPa [27]. Overall, our reported iris stiffness values are consistent and on the same order of magnitude as those reported in the literature, which provides a higher degree of validity to the proposed approach.

It also needs to be noted that the computed stiffness in our study was linearly dependent on the active stress. As our objective was to recreate the deformation pattern of the iris, we argue that with higher input active stress the tissue stiffness would be higher to keep the deformation at the same level.

### Lower permeability in PACG patients

The permeability of the iris was lower in PACG subjects indicating that water absorption and exudation may be reduced in the irides of PACG subjects during smooth muscle contraction. Several studies have reported that the inability of the stroma to lose its interstitial fluid could contribute to the risk of angle-closure [13, 14]. Using OCT imaging, Aptel and Denis demonstrated that the volume and cross-sectional area of the iris decreased with pupil dilation [28]. One of the widely accepted reasoning for the change in iris volume is the gain or loss of extracellular water from the spongy iris stroma [14]. Furthermore, angle-closure eyes demonstrate fewer iris crypts compared to eyes with wider angles [29]. It has been hypothesized that irides with a smaller number of crypts are less porous and would therefore allow less AH to flow through. The decrease in permeability in PACG, as reported in our study, suggests that the stromal tissue would not allow the AH to flow rapidly into or out of the tissue during physiological pupil dilation or constriction, and subsequently, the iris may not change its volume during the deformation process. This condition could potentially cause the iris to bulge significantly at the periphery and lead to an acute angle-closure attack. However, it is still unclear whether a change in permeability would be a cause or the consequence of glaucoma, and further studies are warranted.

We obtained average iris permeability values of 0.55±0.2 mm^2^/kPa.s and 0.41±0.2 mm^2^/kPa.s for healthy and PACG irides, respectively. The current literature on iris tissue permeability is limited. In a previous study by our group, we measured the permeability of porcine stroma ex-vivo and reported a value of 0.0513±0.02 mm^2^/kPa.s. Here our optimized permeability values were an order of magnitude higher than theses experimental observations, but this could potentially be explained by differences in methodologies and the fact that ex-vivo tissues may not be fully representative of their in-vivo state.

### A framework to assess iris biomechanics in-vivo for diagnosing PACG

In this study, we have proposed a non-invasive procedure to evaluate the material parameters of the iris in healthy and PACG conditions. This method does not involve any contact procedure, such as tonometry, or any surgical intervention to isolate the tissue. Furthermore, our method can reproduce the actual motion of the iris during various physiological states. As such, it holds promise as a preferred method for tissue characterization.

The characterization of the biomechanical properties of the iris is important as any deviations from homeostasis may indicate a diseased condition [28]. We observed changes in iris stiffness and permeability in PACG subjects which could indicate that these material parameters may be useful markers for early diagnosis.

### Limitations

In this study, several limitations warrant further discussion. First, we treated the active stress in the sphincter region as a constant. We did not consider any electrophysiological and electromechanical coupling models for sphincter smooth muscle cells [35, 36]. With more advanced models, it may be possible to examine the electrical and chemical response of the iris tissue during miosis and mydriasis. Second, each iris (healthy or PACG) may have its own patient-specific activation force. This was not considered for our optimizations, but instead we used an average for all eyes. Nevertheless, our results still indicate a clear biomechanical difference between two groups, independent of such an assumption. Third, we only used a single OCT B-Scan (horizontal plane) to reconstruct the 3D geometry of the iris during constriction. Using multiple B-Scans in the circumferential direction may yield better results; however, such an approach is not yet feasible as one would have to image the 3D iris in real-time as it deforms during constriction with a high acquisition rate. This may be possible with next-generation OCT devices. Fourth, all tissues (stroma, PEL, sphincter) were described with a simple isotropic Neo-Hookean formulation. Although the active stress in the sphincter region was applied in the circumferential direction, we did not define any circumferential fibers. The rationale behind this was to keep the number of unknown parameters low so that we could avoid the ‘non-uniqueness problem’ that is common in many biomechanical applications. Fifth, all tissues were assumed to be hyperelastic due to their large deformations but not viscoelastic. Several studies have suggested that the iris tissue exhibits viscoelastic properties [30, 31, 32]. Therefore, iris material parameters should be time- and rate-dependent, and the iris should exhibit higher stiffness at higher rates of deformation. As miosis and mydriasis occur rapidly at a rate of approximately 4 mm/s [33], a viscoelastic model may be suitable to describe iris biomechanics and this could be considered in future studies. Sixth, we did not consider the residual stresses that could be present in the iris [34]. Residual stresses can influence the local biomechanical behavior by reducing stress concentrations [17]. Finally, we optimized the material parameters for pupil constriction, and thus, in our simulation, only the sphincter region was active. In the future, we aim to evaluate iris biomechanics while considering both the dilation and constriction of the pupil.

## Conclusion

Our study utilized an inverse FE approach to assess the in-vivo stiffness and permeability of the iris directly from OCT imaging during pupil constriction. We found that PACG eyes exhibited higher iris stiffness but lower permeability. If these tools were to be translated clinically, they could aid in the diagnosis of PACG.

## Acknowledgments

This work was supported by the Singapore Ministry of Education Academic Research Funds Tier 1 (R-397-000-294-114 (MJAG)), the Singapore Ministry of Education Tier 2 (R-397-000-280-112, R-397-000-308-112 (MJAG)), and the National Medical Research Council (Grant NMRC/STAR/ 0023/ 2014 (TA)).

